# Psychedelic Concentrations of Nitrous Oxide Reduce Functional Differentiation in Frontoparietal and Somatomotor Cortical Networks

**DOI:** 10.1101/2023.04.26.538483

**Authors:** Rui Dai, Zirui Huang, Tony E. Larkin, Vijay Tarnal, Paul Picton, Phillip E. Vlisides, Ellen Janke, Amy McKinney, Anthony G. Hudetz, Richard E. Harris, George A. Mashour

## Abstract

Despite the longstanding use of nitrous oxide and descriptions of its psychological effects more than a century ago, there is a paucity of neurobiological investigation of associated psychedelic experiences. Identifying the impact of nitrous oxide on functional brain networks would advance understanding and contribute to the growing body of research in psychedelic neuroscience. Based on human resting-state fMRI data acquired before and during the administration of 35% nitrous oxide, we measured the brain’s functional geometry (through analysis of cortical gradients) and temporal dynamics (through analysis of co-activation patterns). Both analyses show that nitrous oxide reduces functional differentiation in frontoparietal and somatomotor networks. Importantly, the subjective psychedelic experience induced by nitrous oxide is inversely correlated with the degree of functional differentiation. Thus, like classical psychedelics acting on 5-HT2 receptors, nitrous oxide flattens the functional geometry of the cortex and disrupts temporal dynamics in association with psychoactive effects.

## Introduction

As the scientific and clinical interest in psychedelics continues to grow, there is a pressing need for a deeper neurobiological understanding. Although there are extensive neuroimaging studies on the neural correlates of classical serotonergic psychedelics—such as lysergic acid diethylamide (LSD), psilocybin, and dimethyltryptamine (DMT) (Carhart-Harris et al., 2012, 2016; Timmermann et al., 2023)—nitrous oxide remains relatively understudied, despite the longstanding recognition of its psychedelic effects (Block et al., 1990; James, 1874).

Previous human studies using electroencephalography and magnetoencephalography have shown that nitrous oxide alters spectral features, functional connectivity, and complexity (Foster & Liley, 2013; John et al., 2001; Pavone et al., 2016; Pelentritou et al., 2020; Ryu et al., 2017; Vrijdag et al., 2021), but primarily at sedative rather than psychedelic concentrations or without assessment of psychedelic phenomenology. Recently, a functional magnetic resonance imaging (fMRI) study demonstrated functional brain network changes associated with psychedelic effects during nitrous oxide exposure, including a decrease of within-network functional connectivity and an increase of between-network functional connectivity (Dai et al., 2023). This is consistent with previous functional neuroimaging investigations of the acute effect of classical psychedelic drug administration, during which a reduction of functional differentiation of large-scale brain networks was observed (Girn et al., 2023; Kwan et al., 2022; McCulloch et al., 2022). However, the region- or network-specific findings of these studies have shown little convergence (Girn et al., 2022; Madsen et al., 2021; Mason et al., 2020; Müller et al., 2018; Preller et al., 2018, 2020; Roseman et al., 2014; Tagliazucchi et al., 2016), likely constrained by methodological limitations (Girn et al., 2023). First, most studies relied on network-based approaches that were limited to univariate or bivariate analyses, without investigating more distributed multivariate dependencies. Second, most studies utilized time-averaged associations and did not address temporal dynamics.

Recent advances in neuroimaging provide two approaches that can improve understanding of the neural correlates of the psychedelic state. The first approach is cortical gradients, which focuses on multiple functions along a continuum and avoids decomposing the brain into discrete parcellations (Huntenburg et al., 2018; Margulies et al., 2016; Murphy et al., 2018). The second approach is the direct investigation of temporal dynamics, which continuously shape and reshape functional configurations, with spatial patterns of functional connectivity evolving over time to complement functional geometry (Huang et al., 2020; Singleton et al., 2022).

In this study, we investigated whether nitrous oxide reduces functional differentiation in the human cortex, as measured by both functional geometry and temporal dynamics. We reanalyzed a neuroimaging dataset of healthy human volunteers, who were assessed by fMRI before and during exposure to psychedelic concentrations of nitrous oxide and who completed a validated altered-states-of-consciousness questionnaire (Studerus et al., 2010) before and after drug exposure. We quantified the changes of neural activity in cortical gradients and co-activations; we also performed correlation analyses to identify the relationship between subjective psychedelic experience and these brain measures.

## Results

### Nitrous oxide reduces functional differentiation in cortex

In this study, we used cortical gradient analysis to investigate the functional brain connectome in a nonlinear diffusion space. We identified the major spatial axes of functional connectivity at the voxel level, based on the functional similarity structure of fMRI data (i.e., gradients). Voxels within each gradient are categorized according to their similarity in activity patterns, with those at one end of the gradient being more similar (less functionally differentiated) than those at the other. Our results showed that the principal gradient ranged from unimodal cortices (e.g., somatomotor and visual networks) to transmodal cortices (e.g., frontoparietal and default-mode networks) (Figure 1A and 1B), which is consistent with previous studies (Bethlehem et al., 2020; Girn et al., 2022; Huang et al., 2023; Margulies et al., 2016).

**Figure 1:**
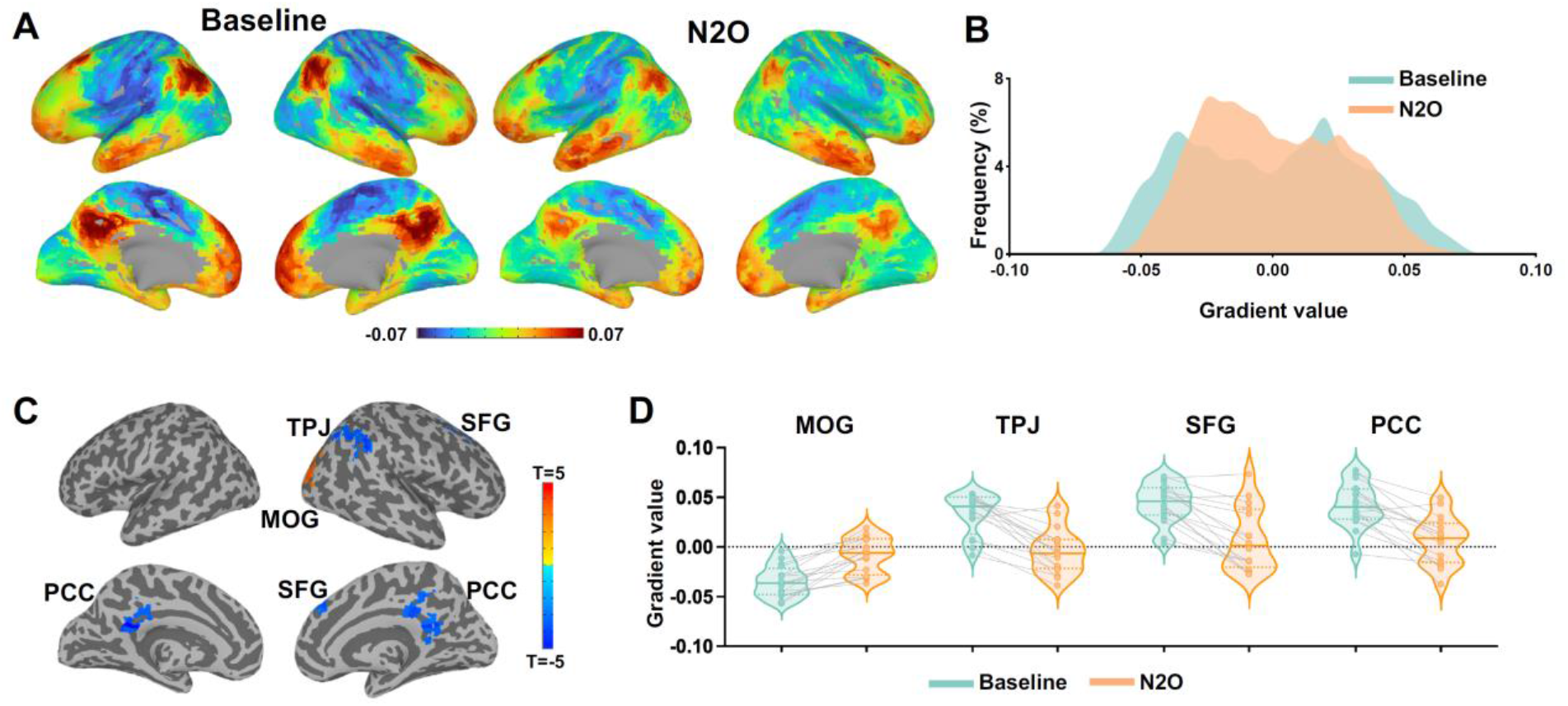
The principal cortical gradient during nitrous oxide and baseline. (A) Visualization of the principal gradient values of the cortex. The color scale represents gradient values, with warm colors indicating transmodal and cool colors indicating unimodal. (B) Histogram of principal gradient derived from whole brain voxels. (C) Voxel-based contrast of the gradient values for nitrous oxide vs. baseline. (D) Individual gradient values extracted from regions showing statistical significance in C.

The voxel-based contrast map (nitrous oxide vs. baseline, pFWE < 0.05 corrected) revealed a decrease of gradient values in the right lateral parietal/temporoparietal junction (TPJ), right superior frontal gyrus (SFG) and bilateral precuneus (PCC), together with an increase of gradient values in the right middle occipital gyrus (MOG), during nitrous oxide administration (Figure 1C and 1D). In addition, we did not find a significant change of the overall range of the principal gradient (Figure S1). Together, the results suggest a center-shifting of specific brain regions along the principal gradient as induced by nitrous oxide, indicating a reduction of differentiation along the functional hierarchy.

We further quantified the cortical gradient changes at the network level. Using a seven-network parcellation scheme (Yeo et al., 2011), we found significant center-shifting during nitrous oxide administration in somatomotor (SMN; t _(15)_ = 6.86, p = 0.00003, Bonferroni-corrected), frontoparietal (FPN; t _(15)_ = 4.98, p = 0.001, Bonferroni-corrected), and default-mode (DMN; t _(15)_ = 3.48, p = 0.02, Bonferroni-corrected) networks (Figure 2). Our network-based results indicated a specific reduction of differentiation at both unimodal end (e.g., SMN) and transmodal end (e.g., FPN and DMN) along the functional hierarchy of the cortex during exposure to psychedelic concentrations of nitrous oxide.

**Figure 2:**
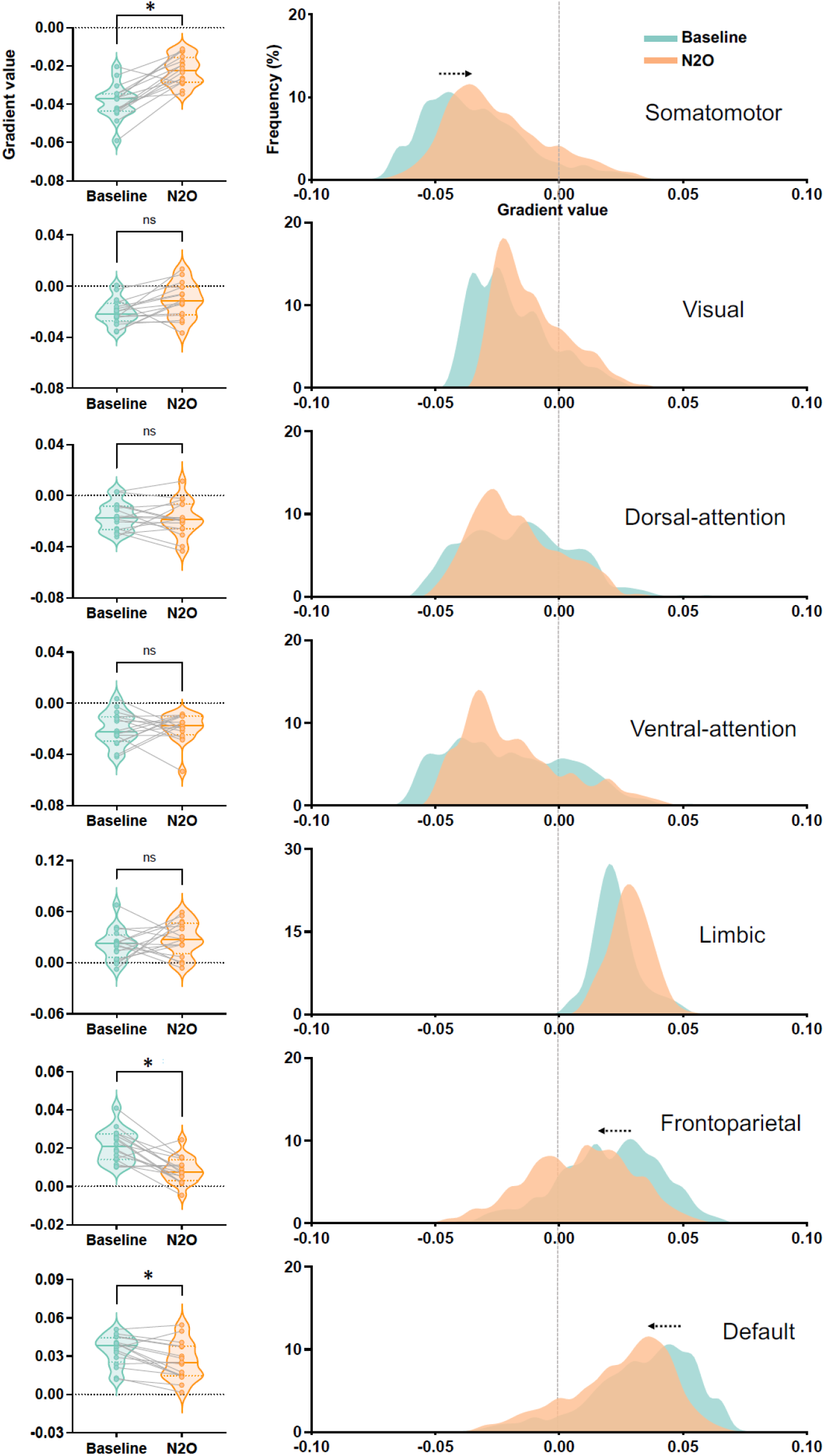
Principal gradient values in different functional networks. Left panel: Paired t-test of gradient values between nitrous oxide and baseline in seven networks. Right panel: histogram of gradient values in nitrous oxide and baseline in seven networks. * Bonferroni-corrected p < 0.05.

Additionally, we conducted an analysis of the second and third gradients of macroscale functional organization. The second gradient showed a range from visual to somatomotor cortices, and the third gradient showed a range from visual and default-mode areas to the areas commonly associated with multiple-demand tasks. However, we did not observe any significant changes during nitrous oxide administration in either gradient (Figure S2 and S3).

### Nitrous oxide disrupts temporal dynamics

In order to examine the temporal evolution of brain activity, we conducted a co-activation pattern (CAP) analysis. Following our previous approach (Huang et al., 2020), we identified eight CAPs, including default mode network (DMN+), dorsal attention network (DAT+), frontoparietal network (FPN+), somatomotor network (SMN+), visual network (VIS+), ventral attention network (VAT+), and global network of activation and deactivation (GN+ and GN−). We calculated the occurrence rates of CAPs by dividing the number of fMRI volumes belonging to a given CAP by the total number of volumes per scan. We compared the occurrence rates between nitrous oxide and baseline in these eight CAPs. Our results revealed a significant reduction in the occurrence rates of FPN+ (t _(15)_ = 4.79, p = 0.002, Bonferroni-corrected) and SMN+ (t _(15)_ = 3.94, p = 0.01, Bonferroni-corrected), accompanied by an increase in the occurrence rates of GN+ (t _(15)_ = 4.62, p = 0.003, Bonferroni-corrected) during nitrous oxide administration. The results indicate that FPN+ and SMN+ occurred less often during nitrous oxide administration, and the two CAPs seemed to be replaced by brain-wide co-activations (i.e., GN+). This is evidence in the temporal domain that nitrous oxide reduces the occurrence of differentiated functional CAPs involving the frontoparietal and somatomotor networks, while promoting a more globally integrated state.

**Figure 3:**
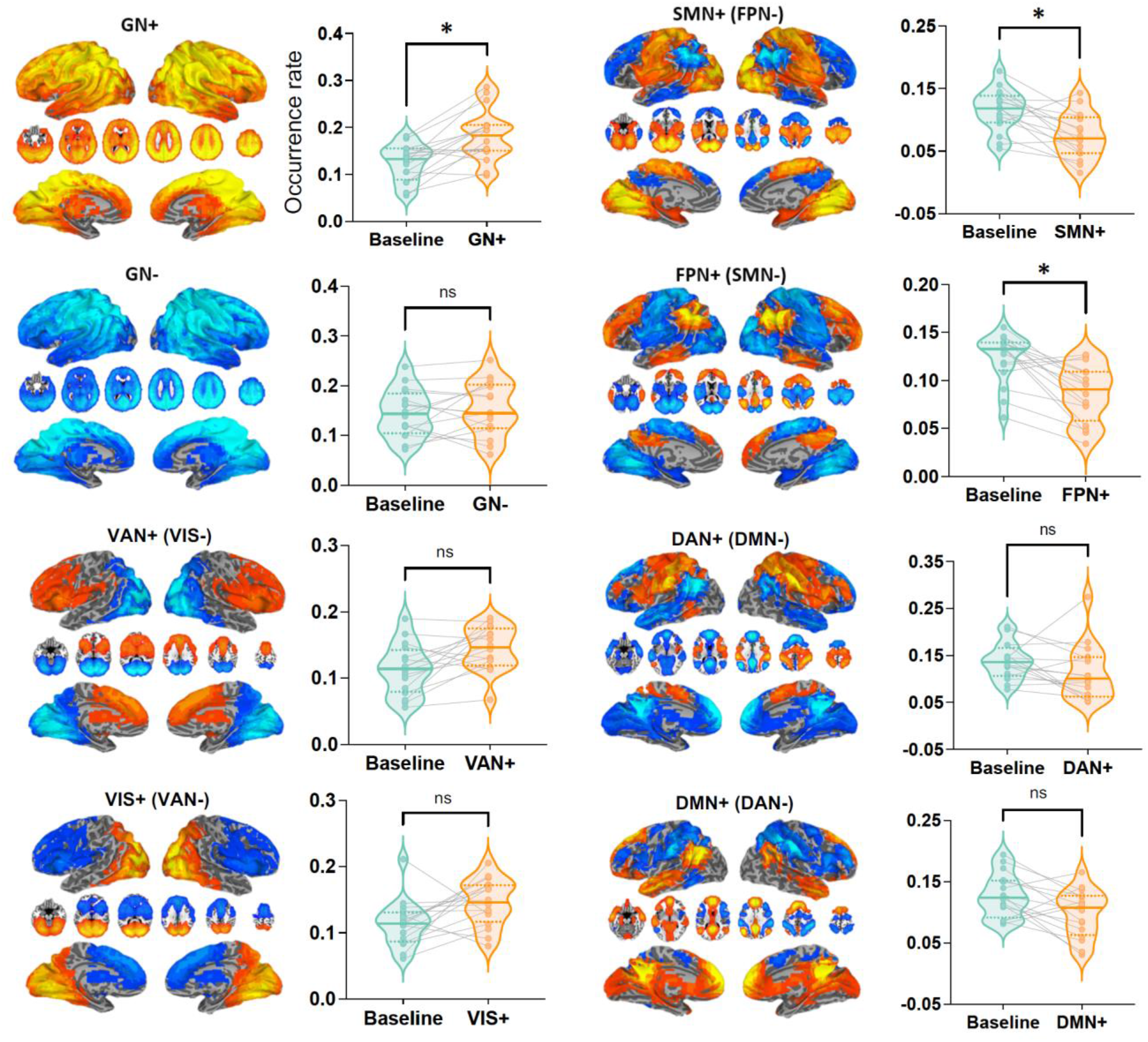
Co-activation patterns and their occurrence rates. Eight co-activation patterns are shown. Paired t-tests of the occurrence rates were performed for nitrous oxide vs. baseline. * Bonferroni-corrected p < 0.05.

In addition, we conducted Spearman correlation analyses to examine the relationship between cortical gradient values and occurrence rates of CAPs in the networks that exhibited significant results in both cortical gradient and CAP analyses (i.e., FPN and SMN). We observed a significant correlation between the FPN gradient score and FPN+ occurrence rate (Figure S4), suggesting that the reduction of functional differentiation in the FPN along the cortical gradient was associated with a decrease in its occurrence rate.

### Psychedelic phenomenology is linked to changes in cortical gradients and dynamic brain activity

To test associations between the degree of changes in cortical gradients and dynamic brain activity with the subjective intensity of the psychedelic state induced by nitrous oxide, we performed correlation analyses between the principal gradient values and the total score derived from altered states of consciousness questionnaire (see Figure S5 for statistics of questionnaire). The analyses involved four regions (MOG, TPJ, SFG, and PCC) identified during voxel-level analysis as well as three networks (SMN, FPN, DMN) identified during network-level analysis. Our results revealed significant correlations between altered states of consciousness and the degree of reduction of functional differentiation in TPJ, PCC, SMN, and FPN (Bonferroni-corrected p < 0.05, Figure 4). These findings suggest that the degree of subjective intensity of the psychedelic state induced by nitrous oxide was associated with changes in cortical network gradients.

**Figure 4:**
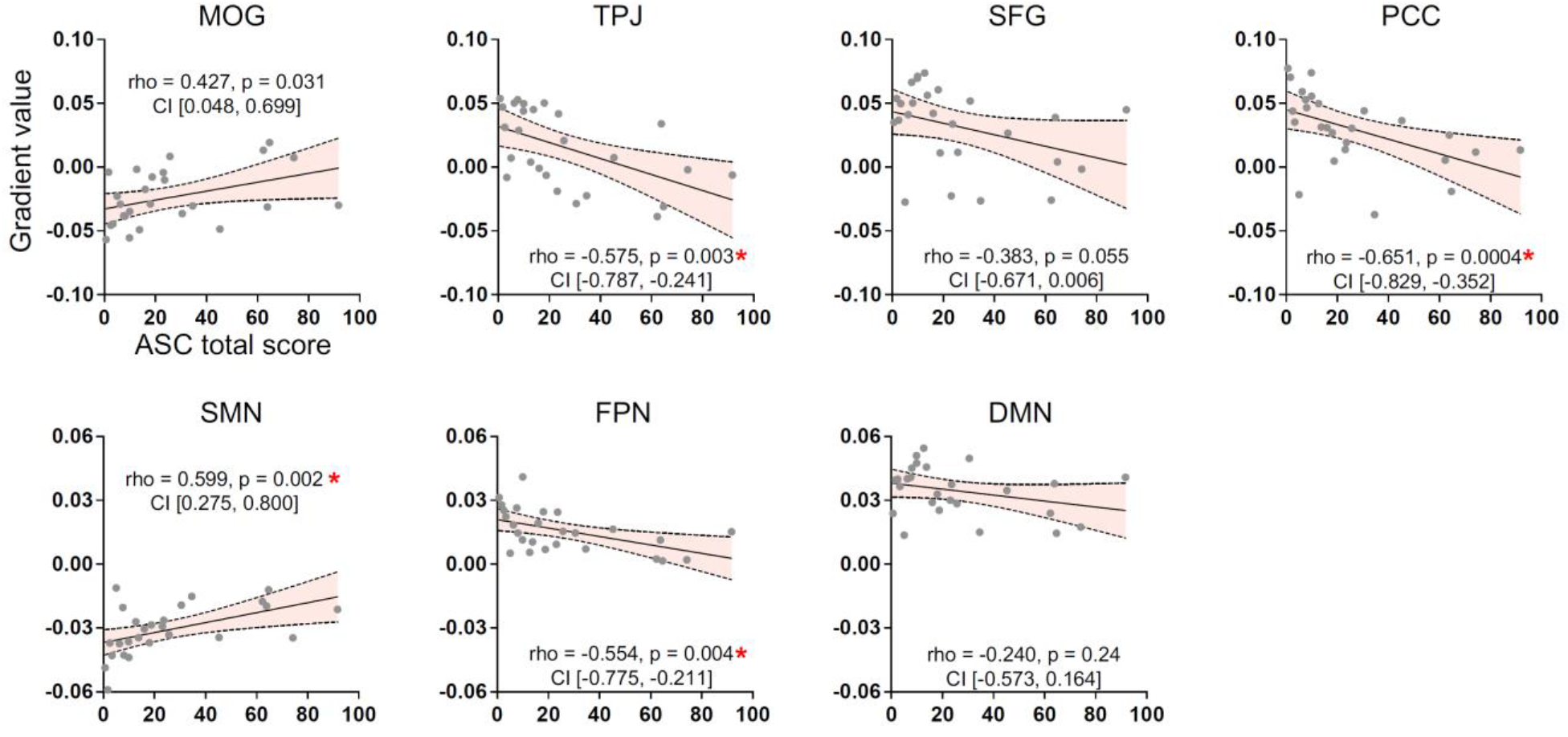
Spearman correlations between gradient values and the total score of 11D-altered states questionnaire. ASC total score measures the intensity of psychedelic experiences, indicating the degree to which an individual’s consciousness deviates from their normal state of consciousness during a psychedelic experience. Higher scores indicate more intense experiences, while lower scores suggest milder effects. Spearman’s rank correlation coefficient (rho), uncorrected p values, and 95% confidence interval (CI) are reported in each scatter plot. * Bonferroni-corrected p < 0.05.

We also conducted correlation analyses between the occurrence rates of CAPs and the total score of altered states of consciousness questionnaire in three networks that displayed statistical significance in the CAP analysis (GN+, SMN+, FPN+). We observed a significant negative correlation in both SMN+ and FPN+ (Figure 5), suggesting that the less frequently these two CAPs occurred, the more intense the psychedelic experience was.

**Figure 5:**
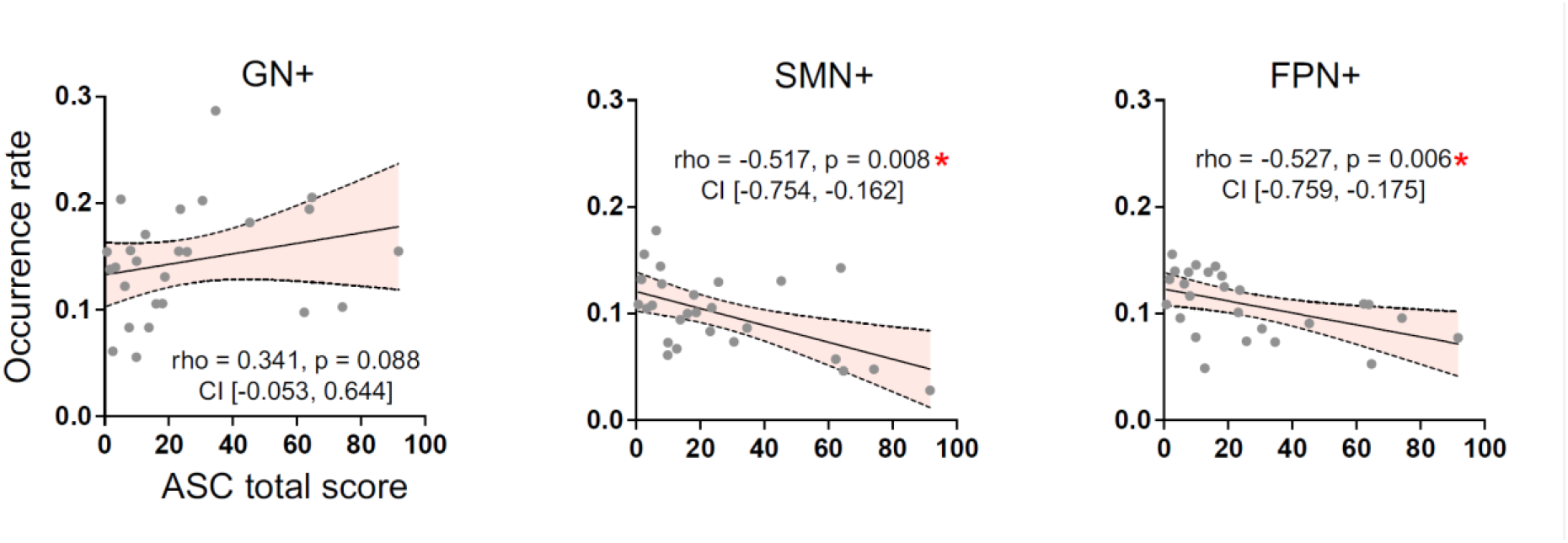
Spearman correlations between gradient values and occurrence rates. Spearman’s rank correlation coefficient (rho), uncorrected p values, and 95% confidence interval (CI) are reported in each scatter plot. * Bonferroni-corrected p < 0.05.

## Discussion

We present evidence that psychedelic concentrations of nitrous oxide induced specific functional reorganizations in the brain, including a reduction of functional differentiation in four brain regions (TPJ, PCC, SFG, and MOG) and three networks (SMN, FPN, and DMN) that were distributed along the hierarchical extremes of the principal/first gradient. Co-activation pattern (CAP) analysis showed a reduced occurrence rate of frontoparietal and somatomotor networks and an enhanced occurrence rate of global co-activation. Finally, we found correlations between the subjective psychedelic experiences and this spatiotemporal reorganization of brain activity. In summary, we present evidence that nitrous oxide flattens the functional geometry of the cortex and disrupts the related temporal dynamics, particularly in frontoparietal and somatomotor networks.

Although most neuroimaging studies have focused on the neural correlates of classical psychedelics such as LSD, psilocybin, and DMT, our study is among the few that investigated nitrous oxide. We utilized two advanced approaches, cortical gradient and CAP analyses, to provide spatially/regionally specific evidence about the neural correlates of the psychedelic state induced by nitrous oxide. A main finding is the reduced functional differentiation in both functional geometry and temporal dynamics, which is aligned with prior research demonstrating that psychedelics reduce functional integration within networks and enhance functional integration between networks (Dai et al., 2023; Girn et al., 2023; Kwan et al., 2022; McCulloch et al., 2022).

More specifically, we found frontoparietal and somatomotor networks were less differentiated relative to the rest of the brain and appeared less frequently in the temporal dynamics; the occurrence of these networks tended to be replaced by increased frequency of global co-active networks. This finding is consistent with previous research demonstrating altered functional connectivity in these networks with classical psychedelics. For example, previous studies of LSD have shown increased global functional connectivity in regions spanning the default mode and frontoparietal networks (Girn et al., 2022; Tagliazucchi et al., 2016) as well as in the somatomotor and visual networks (Preller et al., 2018, 2020). Additionally, recent research has indicated a decrease in functional differentiation at both ends of the hierarchy, encompassing the default and frontoparietal networks at one end and the somatomotor networks at the other end (Girn et al., 2022). Our findings extend these results to the psychedelic state induced by nitrous oxide.

Furthermore, a previous study showed that LSD and psilocybin can flatten the brain’s control energy landscape, which reduces energetic barriers and enables the brain to more easily navigate its repertoire through state transitions (Singleton et al., 2022). In other words, brain states are less “sticky” and can more flexibly move from one state to another (Girn et al., 2023). Our results on nitrous oxide indicate that the flattened energy landscape may be associated, in part, with dedifferentiation of executive control (frontoparietal) and motor control (somatomotor) networks, promoting greater integration and communication between these control systems and other brain areas. However, it is still unclear how a change of functional differentiation along cortical gradients relates to a change of brain entropy (Carhart-Harris, 2018; Carhart-Harris et al., 2014), metastability (Cavanna et al., 2018; Deco & Kringelbach, 2016; Naik et al., 2017), or dynamic repertoires (Hudetz et al., 2015).

We also observed a reduction of functional differentiation in the TPJ and PCC. This finding is in agreement with previous studies that have reported increased global connectivity in these regions under LSD (Tagliazucchi et al., 2016) and increased connectivity between the right TPJ and other regions of the posterior cortex during exposure to nitrous oxide, ketamine, and LSD (Dai et al., 2023). Together, our results suggest that the modulation of these regions may contribute to the phenomenology of the psychedelic experience (Vlisides et al., 2018).

There are numerous limitations to this investigation. First, our results were primarily based on the cortex; future investigation of subcortical regions using high-field MRI will be important to clarify fully the neurobiology of the psychedelic state induced by nitrous oxide. Second, the present study exclusively examined the neural correlates associated with nitrous oxide, highlighting the need for comparison with other psychoactive drugs such as ketamine, dimethyltryptamine, and methylenedioxymethamphetamine. Despite these limitations, this study is the first to characterize cortical gradient and temporal dynamic changes during the administration of psychedelic doses of nitrous oxide. The present study extends previous findings on the neural underpinning of the psychedelic state induced by 5-HT2 modulators and provides novel neural correlates of altered subjective experiences induced by nitrous oxide.

## Methods

The study was performed at the University of Michigan Medical School and received approval from the Institutional Review Board under the identifier HUM00096321. Prior to participation, all subjects were carefully informed about the potential risks and benefits of the study, and written informed consent was obtained from each individual. The analysis of this study was part of a clinical trial that was registered with clinicaltrials.gov under the identifier NCT03435055, and the primary study’s results were released in July 2021.

### Participants

In this study, sixteen healthy participants (8 males, means ± SD, ages: 24.6±3.7 years) underwent two resting-state fMRI scans before and during exposure to subanesthetic levels of nitrous oxide (i.e., 35% concentration). Two participants were excluded because of excessive head motion (more than a half TRs in each scan). Requirements for participation, included being classified as physical status I by the American Society of Anesthesiologists, being free of drug abuse or psychosis, and being free of other health-related conditions (https://www.clinicaltrials.gov/ct2/show/NCT03435055).

### Experimental Design

The study consisted of two visits for participants; a pre-scan visit and a scanning visit within three days. During the pre-scan visit, participants were informed of the study protocol. During the scanning visit, participants underwent fMRI data collection during both placebo and subanesthetic nitrous oxide inhalation. Prior to the resting state scan, nitrous oxide was administered to achieve at least 5 minutes of equilibrium, and any adverse physiological or psychological reactions were monitored and addressed. Before scanning and after 30 minutes of recovery from nitrous oxide administration, the altered states consciousness questionnaire (Studerus et al., 2010) was administered. Thirteen out of the 16 participants completed the survey.

### Drug Administration

The administration of nitrous oxide was conducted using MRI-compatible anesthesia machines, overseen by at least two fully trained anesthesiologists. Prior to imaging, nitrous oxide was first administered outside of the scanner to ensure airway patency and physiological stability. To mitigate predicable common side effects, participants were given ondansetron (4-8 mg IV) with an additional dose of dexamethasone (4 mg IV) if necessary; glycopyrrolate (0.2-0.4 mg IV), labetalol (5-10 mg/kg IV), and midazolam (1-2 mg IV) were available if necessary. Standard intraoperative monitoring devices, including electrocardiogram, blood pressure, pulse oximetry, and capnography, were used throughout the experiment. To reduce interference from external stimuli, participants wore earplugs and headphones during the fMRI scanning.

### fMRI data acquisition

Imaging data were obtained using a 3T Philips Achieva MRI scanner (Best, Netherlands) located at Michigan Medicine, University of Michigan. Functional whole-brain images were acquired using a T2*-weighted echo-planar sequence with the following parameters: 48 slices, TR/TE = 2000/30ms, slice thickness = 3 mm, field of view = 200 × 200mm, flip angle = 90°, and scan time of 6 minutes. High-resolution anatomical images were also acquired for co-registration with the resting state fMRI data.

### Altered States Questionnaire

There are 11 dimensions included in the altered-states-of-consciousness questionnaire, including the following: experiences of unity, spiritual experience, blissful state, insightfulness, disembodiment, impaired control and cognition, anxiety, complex imagery, elementary imagery, audiovisual synesthesia, and changed meaning of percepts. Participants were asked to rate their experiences on each dimension using a discrete response scale with 11 options, ranging from 0 (Never) to 10 (Always). The reported scale scores were calculated by averaging the responses across all items belonging to each respective scale.

### fMRI data preprocessing

The fMRI data preprocessing steps were conducted using AFNI (http://afni.nimh.nih.gov/). The following procedures were applied: (1) Removal of the first two frames of each scan; (2) Slice timing correction; (3) Rigid head motion correction/realignment. The frame-wise displacement (FD) of head motion was calculated as the Euclidean Norm of the six-dimension motion derivatives. Any frame with a derivative value exceeding the FD of 0.4mm was removed, along with its previous frame; (4) Coregistration with T1 anatomical images; (5) Spatial normalization into Talaraich stereotactic space and resampling to 4 mm isotropic voxels; (6) Time-censored data was band-pass filtered to 0.01–0.1Hz using AFNI’s function 3dTproject. Linear regression was used to remove undesired components such as linear and nonlinear drift, time series of head motion and its temporal derivative, and mean time series from the white matter and cerebrospinal fluid; (7) Spatial smoothing with a 6mm full-width at half-maximum isotropic Gaussian kernel; (8) Normalization of each voxel’s time series to zero mean and unit variance.

### Cortical gradient analysis

Cortical gradient analysis was performed at the voxel level. The fMRI time series were first extracted from each voxel, and a 14,871 × 14,871 connectivity matrix was constructed for each participant and condition using Pearson correlation. To obtain a group-average connectivity matrix for each condition, individual matrices were averaged. The Brain Space toolbox (https://brainspace.readthedocs.io/en/ latest/) implemented in MATLAB R2022a was utilized for conducting cortical gradients analysis (Vos de Wael et al., 2020). Based on previous work (Margulies et al., 2016; Mckeown et al., 2020; Raut et al., 2021), the connectivity matrix was first z-transformed and thresholded at a sparsity of 90%. This resulted in only the top 10% of weighted connections per row being retained. Subsequently, a normalized cosine angle affinity matrix was calculated to measure the similarity of connectivity profiles between different cortical areas. Based on a diffusion map embedding algorithm, gradient components were identified, which estimated the low-dimensional embedding from the high-dimensional connectivity matrix The parameter α determines the degree to which the density of sampling points on the manifold influences the algorithm, where α=0 signifies maximal influence and α=1 indicates no influence. Additionally, the parameter t controls the scale of eigenvalues of the diffusion operator. Following recommendations of previous studies (Bethlehem et al., 2020; Hong et al., 2019; Margulies et al., 2016; Mckeown et al., 2020; Vos de Wael et al., 2020), we fixed global relations between data points in the embedded space by setting α to 0.5 and t to 0, which indicated that the estimation of diffusion time is automated and derived through a damped regularization process. (Margulies et al., 2016; Vos de Wael et al., 2020). At the group-level, the gradient solutions were aligned to a subsample of the human connectome project dataset (n=100) using Procrustes rotation. This method involves finding an orthogonal linear transformation that superimposes a given source S onto a target T representation, effectively aligning the two representations (Langs et al., 2014). The Procrustes rotation transformation was implemented to address the issue of eigenvector multiplicity and sign ambiguity, which may cause the computed gradients from different individuals to be incomparable. The alignment step improves the stability of gradient estimation and enables better comparability with existing literature (Vos de Wael et al., 2020). Individual-level gradients for each condition were computed using the same set of parameters. The gradient eigenvector loading values from seven pre-defined functional networks (Yeo et al., 2011) were used to depict the cortical gradient organization at the network level.

### Tracking large-scale co-activation patterns

A spatial similarity analysis was performed to evaluate the relationship between the signal intensity of each fMRI volume and eight pre-defined centroids of co-activation patterns (CAPs) described by our previous study (Huang et al., 2020). The CAPs consisted of the DMN+, DAN+, FPN+, SMN+, VIS+, VAN+, and global network of activation and deactivation (GN+ and GN−). Each fMRI volume was assigned to a specific CAP based on its maximal similarity to the CAP centroids. This enabled us to generate a time series of discrete CAP labels. We then computed the occurrence rate of each CAP at the individual level by dividing the number of volumes assigned to a specific CAP by the total number of volumes per state or condition.

### Statistical analysis

To correct for multiple comparisons in voxel-based gradient analysis, Monte Carlo simulation was used via the AFNI program 3dClustSim, resulting in a family-wise error rate of p < 0.05 with a minimum cluster size of 60 voxels. For network-based gradient analysis (mean gradient values) and CAP analysis, paired t-tests were conducted comparing the nitrous oxide condition to the baseline. The Spearman correlations were used to analyze the relationship between the altered states questionnaire total score and gradient value, as well as between the altered states questionnaire total score and CAP occurrence rate. Bonferroni correction, i.e., dividing the critical P value α=0.05 by the number of comparisons being made, was used to counteract the multiple comparisons problem. JASP (v0.16.3; https://jasp-stats.org/) was used for statistical calculations.

## Data and code availability statement

Publicly available software used for analyses is AFNI and Matlab. All data needed to evaluate the conclusions in this article are present in the main text and the Supplementary Materials. Access to additional data by qualified investigators (i.e., affiliated with accredited academic and research institutions) are subject to scientific and ethical review. Completion of a material transfer agreement signed by an institutional official will be required in order to access the data.

## Ethics statement

This study was conducted at the University of Michigan Medical School, where Institutional Review Board (IRB, HUM00096321) approval was obtained. The study team carefully discussed risks and benefits with all participants, after which written informed consent was documented. This analysis was part of a clinical study registered with clinicaltrials.gov (NCT03435055); results from the primary study were posted in July 2021. All methods were performed in accordance with the relevant guidelines and regulations and written informed consent was obtained.

## Supplementary Material

**Figure S1:**
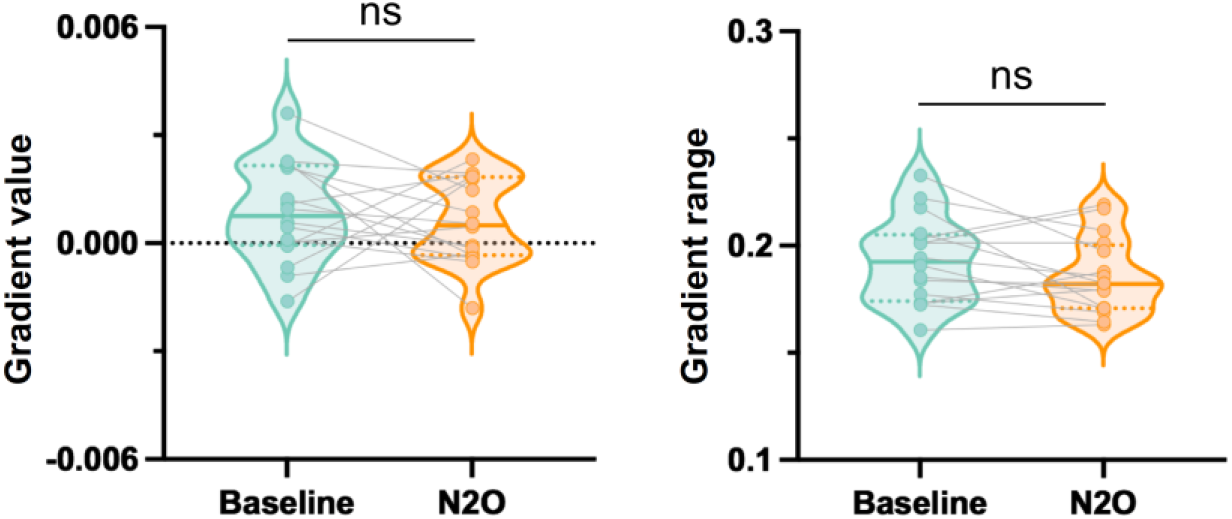
Principal functional cortical gradient value and range during nitrous oxide and baseline. Left panel: Paired t-test of the gradient value between nitrous oxide and baseline. Right panel: Paired t-test of the gradient range between nitrous oxide and baseline. Numerical range of each gradient was calculated as the distance from the minimum to the maximum gradient eigenvector values, indicating segregation (i.e., different connectivity profile) of the gradient extremes.

**Figure S2:**
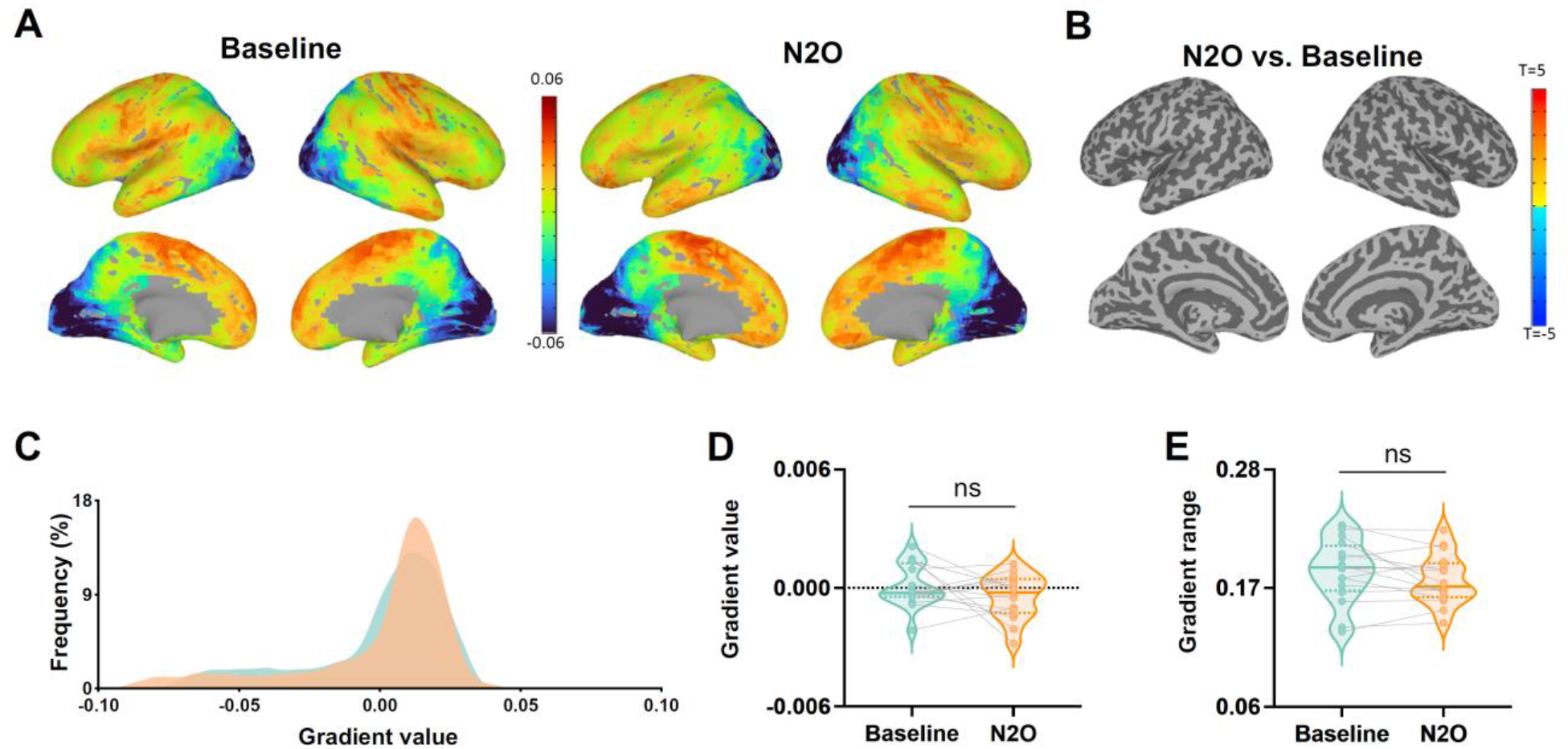
Secondary functional cortical gradient during nitrous oxide and baseline. (A) Global gradient values from visual to somatomotor axis. (B) Voxel-based contrast of the gradient values by nitrous oxide vs. baseline. (C) Global histogram of the secondary gradient. (D) Paired t-test of the gradient value between nitrous oxide and baseline. (E) Paired t-test of the gradient range between nitrous oxide and baseline.

**Figure S3:**
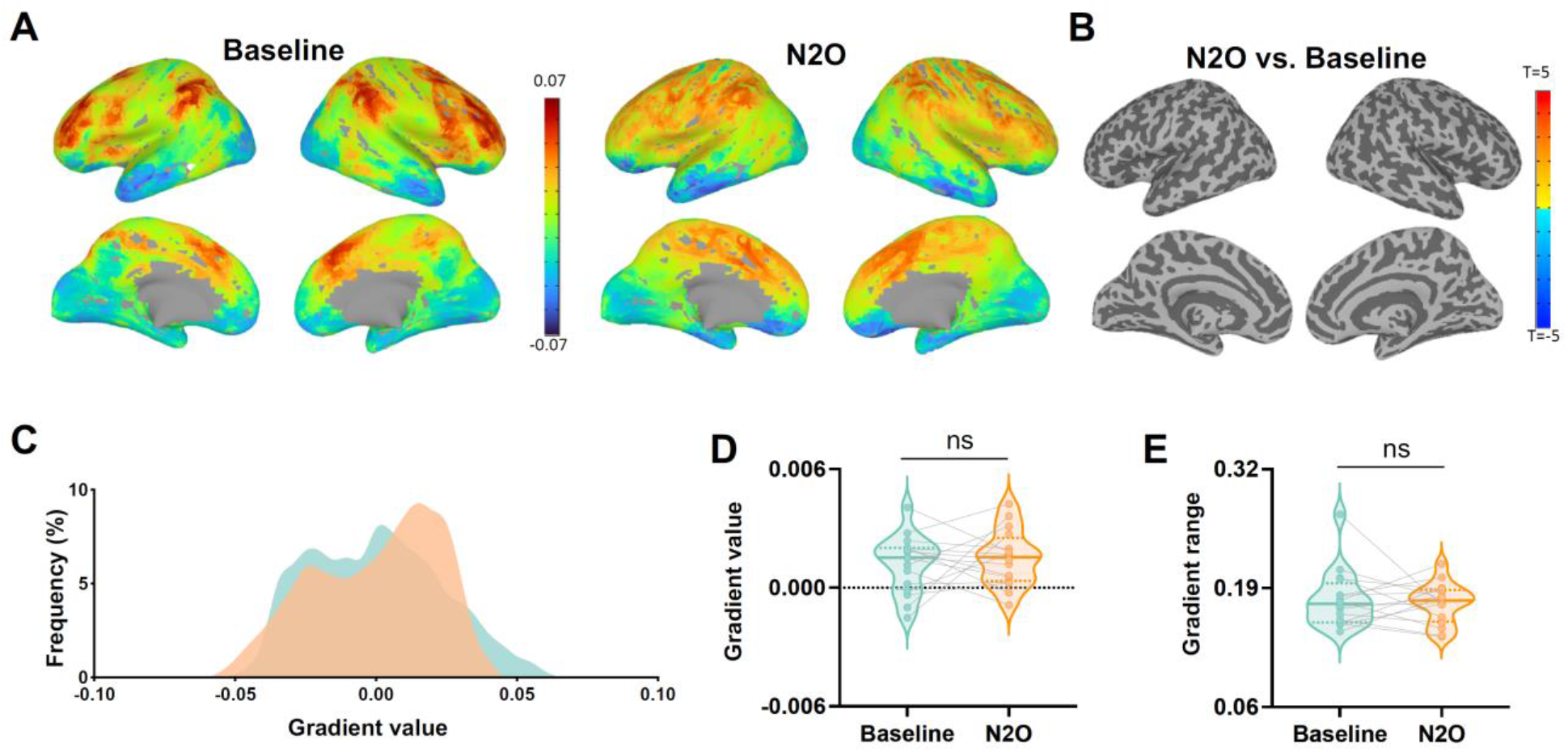
Third functional cortical gradient during nitrous oxide and baseline. (A) Global gradient values from visual to default-mode axis. (B) Voxel-based contrast of the gradient values by nitrous oxide vs. baseline. (C) Global histogram of the third gradient. (D) Paired t-test of the gradient value between nitrous oxide and baseline. (E) Paired t-test of the gradient range between nitrous oxide and baseline.

**Figure S4:**
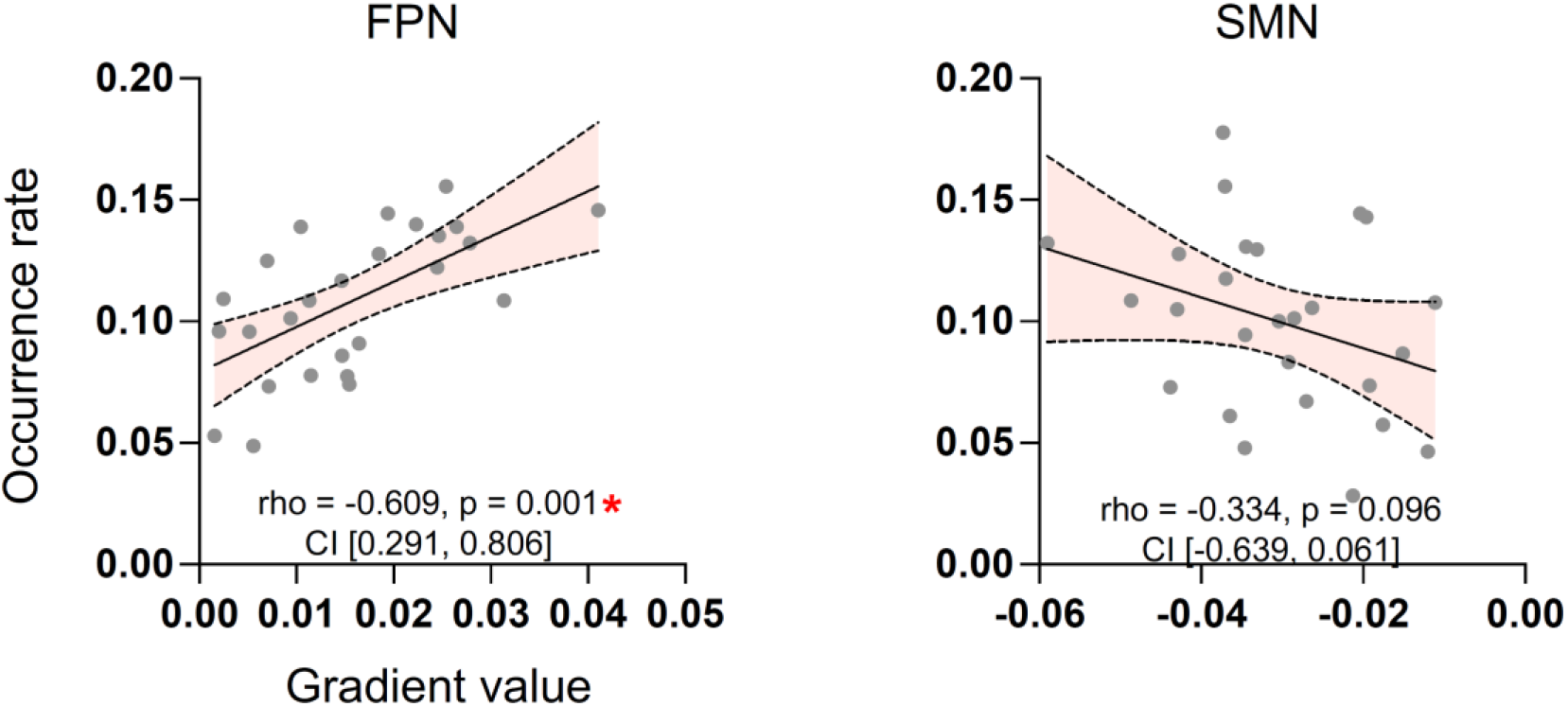
Spearman correlations between gradient score and occurrence rate. Spearman’s rank correlation coefficient (rho), uncorrected p values, and 95% confidence interval (CI) are reported in each scatter plot. * Bonferroni-corrected p < 0.05.

**Figure S5:**
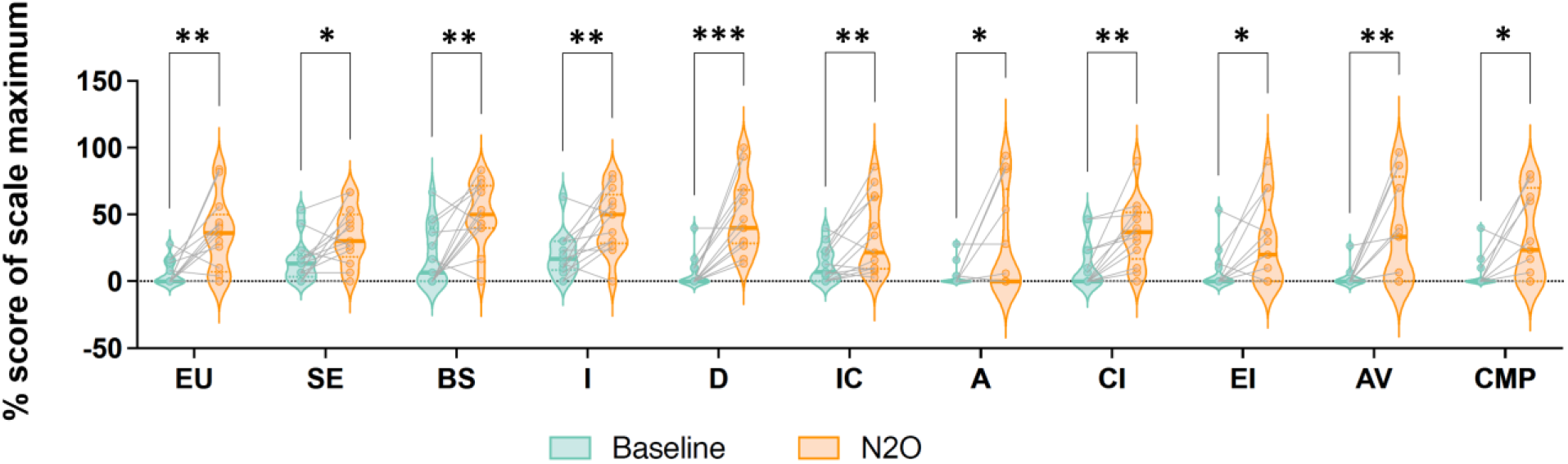
Behavioral results derived from the 11D-altered states questionnaire. Nitrous oxide (N2O) administration resulted in significantly higher total scores than that of baseline condition (mean ± SEM, baseline: 10.01 ± 2.88, nitrous oxide: 37.12 ± 6.69, t (12) = 3.86, p = 0.002). Subscales include the following, EU: experience of unity, SE: spiritual experience, BS: blissful state, I: insightfulness, D: disembodiment, IC: impaired control and cognition, A: anxiety, CI: complex imagery, EI: elementary imagery, AV: audio-visual synesthesia, CMP: changed meaning of percepts. * p < 0.05, ** p < 0.01, *** p < 0.001.

## References

Bethlehem, R. A. I., Paquola, C., Seidlitz, J., Ronan, L., Bernhardt, B., Consortium, C. C. A. N., & Tsvetanov, K. A. (2020). Dispersion of functional gradients across the adult lifespan. NeuroImage, 222(July), 117299. https://doi.org/10.1016/j.neuroimage.2020.117299

Block, R. I., Ghoneim, M. M., Kumar, V., & Pathak, D. (1990). Psychedelic effects of a subanesthetic concentration of nitrous oxide. Anesthesia Progress, 37(6), 271–276. http://www.ncbi.nlm.nih.gov/pubmed/2097905

Carhart-Harris, R. L. (2018). The entropic brain - revisited. Neuropharmacology, 142, 167–178. https://doi.org/10.1016/j.neuropharm.2018.03.010

Carhart-Harris, R. L., Erritzoe, D., Williams, T., Stone, J. M., Reed, L. J., Colasanti, A., Tyacke, R. J., Leech, R., Malizia, A. L., Murphy, K., Hobden, P., Evans, J., Feilding, A., Wise, R. G., & Nutt, D. J. (2012). Neural correlates of the psychedelic state as determined by fMRI studies with psilocybin. Proceedings of the National Academy of Sciences of the United States of America, 109(6), 2138– 2143. https://doi.org/10.1073/pnas.1119598109

Carhart-Harris, R. L., Leech, R., Hellyer, P. J., Shanahan, M., Feilding, A., Tagliazucchi, E., Chialvo, D. R., & Nutt, D. (2014). The entropic brain: A theory of conscious states informed by neuroimaging research with psychedelic drugs. Frontiers in Human Neuroscience, 8(1 FEB), 1–22. https://doi.org/10.3389/fnhum.2014.00020

Carhart-Harris, R. L., Muthukumaraswamy, S., Roseman, L., Kaelen, M., Droog, W., Murphy, K., Tagliazucchi, E., Schenberg, E. E., Nest, T., Orban, C., Leech, R., Williams, L. T., Williams, T. M., Bolstridge, M., Sessa, B., McGonigle, J., Sereno, M. I., Nichols, D., Hellyer, P. J., … Nutt, D. J. (2016). Neural correlates of the LSD experience revealed by multimodal neuroimaging. Proceedings of the National Academy of Sciences of the United States of America, 113(17), 4853–4858. https://doi.org/10.1073/pnas.1518377113

Cavanna, F., Vilas, M. G., Palmucci, M., & Tagliazucchi, E. (2018). Dynamic functional connectivity and brain metastability during altered states of consciousness. NeuroImage, 180, 383–395. https://doi.org/10.1016/j.neuroimage.2017.09.065

Dai, R., Larkin, T. E., Huang, Z., Tarnal, V., Picton, P., Vlisides, P. E., Janke, E., McKinney, A., Hudetz, A. G., Harris, R. E., & Mashour, G. A. (2023). Classical and Non-Classical Psychedelic Drugs Induce Common Network Changes in Human Cortex. NeuroImage, 120097. https://doi.org/10.1016/j.neuroimage.2023.120097

Deco, G., & Kringelbach, M. L. (2016). Metastability and Coherence: Extending the Communication through Coherence Hypothesis Using A Whole-Brain Computational Perspective. Trends in Neurosciences, 39(3), 125–135. https://doi.org/10.1016/j.tins.2016.01.001

Foster, B. L., & Liley, D. T. J. (2013). Effects of nitrous oxide sedation on resting electroencephalogram topography. Clinical Neurophysiology: Official Journal of the International Federation of Clinical Neurophysiology, 124(2), 417–423. https://doi.org/10.1016/j.clinph.2012.08.007

Girn, M., Rosas, F. E., Daws, R. E., Gallen, C. L., Gazzaley, A., & Carhart-Harris, R. L. (2023). A complex systems perspective on psychedelic brain action. Trends in Cognitive Sciences, 1–13. https://doi.org/10.1016/j.tics.2023.01.003

Girn, M., Roseman, L., Bernhardt, B., Smallwood, J., Carhart-Harris, R., & Nathan Spreng, R. (2022). Serotonergic psychedelic drugs LSD and psilocybin reduce the hierarchical differentiation of unimodal and transmodal cortex. NeuroImage, 256(April), 119220. https://doi.org/10.1016/j.neuroimage.2022.119220

Hong, S. J., de Wael, R. V., Bethlehem, R. A. I., Lariviere, S., Paquola, C., Valk, S. L., Milham, M. P., Di Martino, A., Margulies, D. S., Smallwood, J., & Bernhardt, B. C. (2019). Atypical functional connectome hierarchy in autism. Nature Communications, 10(1), 1–13. https://doi.org/10.1038/s41467-019-08944-1

Huang, Z., Mashour, G. A., & Hudetz, A. G. (2023). Functional geometry of the cortex encodes dimensions of consciousness. Nature Communications, 14(1), 72. https://doi.org/10.1038/s41467-022-35764-7

Huang, Z., Zhang, J., Wu, J., Mashour, G. A., & Hudetz, A. G. (2020). Temporal circuit of macroscale dynamic brain activity supports human consciousness. Science Advances, 6(11), 1–15. https://doi.org/10.1126/sciadv.aaz0087

Hudetz, A. G., Liu, X., & Pillay, S. (2015). Dynamic repertoire of intrinsic brain states is reduced in propofol-induced unconsciousness. Brain Connectivity, 5(1), 10–22. https://doi.org/10.1089/brain.2014.0230

Huntenburg, J. M., Bazin, P. L., & Margulies, D. S. (2018). Large-Scale Gradients in Human Cortical Organization. Trends in Cognitive Sciences, 22(1), 21–31. https://doi.org/10.1016/j.tics.2017.11.002

James, W. (1874). Review of “The Anaesthetic Revelation and the Gist of Philosophy.” The Atlantic Monthly, 33(205), 627–628.

John, E. R., Prichep, L. S., Kox, W., Valdés-Sosa, P., Bosch-Bayard, J., Aubert, E., Tom, M., di Michele, F., Gugino, L. D., & DiMichele, F. (2001). Invariant reversible QEEG effects of anesthetics. Consciousness and Cognition, 10(2), 165–183. https://doi.org/10.1006/ccog.2001.0507

Kwan, A. C., Olson, D. E., Preller, K. H., & Roth, B. L. (2022). The neural basis of psychedelic action. 25(November). https://doi.org/10.1038/s41593-022-01177-4

Langs, G., Sweet, A., Lashkari, D., Tie, Y., Rigolo, L., Golby, A. J., & Golland, P. (2014). Decoupling function and anatomy in atlases of functional connectivity patterns: Language mapping in tumor patients. NeuroImage, 103, 462–475. https://doi.org/10.1016/j.neuroimage.2014.08.029

Madsen, M. K., Stenbæk, D. S., Arvidsson, A., Armand, S., Marstrand-Joergensen, M. R., Johansen, S. S., Linnet, K., Ozenne, B., Knudsen, G. M., & Fisher, P. M. (2021). Psilocybin-induced changes in brain network integrity and segregation correlate with plasma psilocin level and psychedelic experience. European Neuropsychopharmacology: The Journal of the European College of Neuropsychopharmacology, 50, 121–132. https://doi.org/10.1016/j.euroneuro.2021.06.001

Margulies, D. S., Ghosh, S. S., Goulas, A., Falkiewicz, M., Huntenburg, J. M., Langs, G., Bezgin, G., Eickhoff, S. B., Castellanos, F. X., Petrides, M., Jefferies, E., & Smallwood, J. (2016). Situating the default-mode network along a principal gradient of macroscale cortical organization. Proceedings of the National Academy of Sciences of the United States of America, 113(44), 12574–12579. https://doi.org/10.1073/pnas.1608282113

Mason, N. L., Kuypers, K. P. C., Müller, F., Reckweg, J., Tse, D. H. Y., Toennes, S. W., Hutten, N. R. P. W., Jansen, J. F. A., Stiers, P., Feilding, A., & Ramaekers, J. G. (2020). Me, myself, bye: regional alterations in glutamate and the experience of ego dissolution with psilocybin. Neuropsychopharmacology: Official Publication of the American College of Neuropsychopharmacology, 45(12), 2003–2011. https://doi.org/10.1038/s41386-020-0718-8

McCulloch, D. E. W., Knudsen, G. M., Barrett, F. S., Doss, M. K., Carhart-Harris, R. L., Rosas, F. E., Deco, G., Kringelbach, M. L., Preller, K. H., Ramaekers, J. G., Mason, N. L., Müller, F., & Fisher, P. M. D. (2022). Psychedelic resting-state neuroimaging: A review and perspective on balancing replication and novel analyses. Neuroscience and Biobehavioral Reviews, 138. https://doi.org/10.1016/j.neubiorev.2022.104689

Mckeown, B., Strawson, W. H., Wang, H. T., Karapanagiotidis, T., Vos de Wael, R., Benkarim, O., Turnbull, A., Margulies, D., Jefferies, E., McCall, C., Bernhardt, B., & Smallwood, J. (2020). The relationship between individual variation in macroscale functional gradients and distinct aspects of ongoing thought. NeuroImage, 220(May). https://doi.org/10.1016/j.neuroimage.2020.117072

Müller, F., Dolder, P. C., Schmidt, A., Liechti, M. E., & Borgwardt, S. (2018). Altered network hub connectivity after acute LSD administration. NeuroImage: Clinical, 18, 694–701. https://doi.org/10.1016/j.nicl.2018.03.005

Murphy, C., Jefferies, E., Rueschemeyer, S.-A., Sormaz, M., Wang, H.-T., Margulies, D. S., & Smallwood, J. (2018). Distant from input: Evidence of regions within the default mode network supporting perceptually-decoupled and conceptually-guided cognition. NeuroImage, 171(January), 393–401. https://doi.org/10.1016/j.neuroimage.2018.01.017

Naik, S., Banerjee, A., Bapi, R. S., Deco, G., & Roy, D. (2017). Metastability in Senescence. Trends in Cognitive Sciences, 21(7), 509–521. https://doi.org/10.1016/j.tics.2017.04.007

Pavone, K. J., Akeju, O., Sampson, A. L., Ling, K., Purdon, P. L., & Brown, E. N. (2016). Nitrous oxide-induced slow and delta oscillations. Clinical Neurophysiology: Official Journal of the International Federation of Clinical Neurophysiology, 127(1), 556–564. https://doi.org/10.1016/j.clinph.2015.06.001

Pelentritou, A., Kuhlmann, L., Cormack, J., Mcguigan, S., Woods, W., Muthukumaraswamy, S., & Liley, D. (2020). Source-level Cortical Power Changes for Xenon and Nitrous Oxide-induced Reductions in Consciousness in Healthy Male Volunteers. Anesthesiology, 132(5), 1017–1033. https://doi.org/10.1097/ALN.0000000000003169

Preller, K. H., Burt, J. B., Ji, J. L., Schleifer, C. H., Adkinson, B. D., Stämpfli, P., Seifritz, E., Repovs, G., Krystal, J. H., Murray, J. D., Vollenweider, F. X., & Anticevic, A. (2018). Changes in global and thalamic brain connectivity in LSD-induced altered states of consciousness are attributable to the 5-HT2A receptor. ELife, 7. https://doi.org/10.7554/eLife.35082

Preller, K. H., Duerler, P., Burt, J. B., Ji, J. L., Adkinson, B., Stämpfli, P., Seifritz, E., Repovš, G., Krystal, J. H., Murray, J. D., Anticevic, A., & Vollenweider, F. X. (2020). Psilocybin Induces Time-Dependent Changes in Global Functional Connectivity. Biological Psychiatry, 88(2), 197–207. https://doi.org/10.1016/j.biopsych.2019.12.027

Raut, R. V., Snyder, A. Z., Mitra, A., Yellin, D., Fujii, N., Malach, R., & Raichle, M. E. (2021). Global waves synchronize the brain’s functional systems with fluctuating arousal. Science Advances, 7(30), 1–16. https://doi.org/10.1126/sciadv.abf2709

Roseman, L., Leech, R., Feilding, A., Nutt, D. J., & Carhart-Harris, R. L. (2014). The effects of psilocybin and MDMA on between-network resting state functional connectivity in healthy volunteers. Frontiers in Human Neuroscience, 8(MAY), 1–11. https://doi.org/10.3389/fnhum.2014.00204

Ryu, J.-H., Kim, P.-J., Kim, H.-G., Koo, Y.-S., & Shin, T. J. (2017). Investigating the effects of nitrous oxide sedation on frontal-parietal interactions. Neuroscience Letters, 651, 9–15. https://doi.org/10.1016/j.neulet.2017.04.036

Singleton, S. P., Luppi, A. I., Carhart-Harris, R. L., Cruzat, J., Roseman, L., Nutt, D. J., Deco, G., Kringelbach, M. L., Stamatakis, E. A., & Kuceyeski, A. (2022). Receptor-informed network control theory links LSD and psilocybin to a flattening of the brain’s control energy landscape. Nature Communications, 13(1), 1–13. https://doi.org/10.1038/s41467-022-33578-1

Studerus, E., Gamma, A., & Vollenweider, F. X. (2010). Psychometric evaluation of the altered states of consciousness rating scale (OAV). PloS One, 5(8), e12412. https://doi.org/10.1371/journal.pone.0012412

Tagliazucchi, E., Roseman, L., Kaelen, M., Orban, C., Muthukumaraswamy, S. D., Murphy, K., Laufs, H., Leech, R., McGonigle, J., Crossley, N., Bullmore, E., Williams, T., Bolstridge, M., Feilding, A., Nutt, D. J., & Carhart-Harris, R. (2016). Increased Global Functional Connectivity Correlates with LSD-Induced Ego Dissolution. Current Biology, 26(8), 1043–1050. https://doi.org/10.1016/j.cub.2016.02.010

Timmermann, C., Roseman, L., Haridas, S., Rosas, F. E., Luan, L., Kettner, H., Martell, J., Erritzoe, D., Tagliazucchi, E., Pallavicini, C., Girn, M., Alamia, A., Leech, R., Nutt, D. J., & Carhart-Harris, R. L. (2023). Human brain effects of DMT assessed via EEG-fMRI. Proceedings of the National Academy of Sciences, 120(13), 2017. https://doi.org/10.1073/pnas.2218949120

Vlisides, P. E., Bel-Bahar, T., Nelson, A., Chilton, K., Smith, E., Janke, E., Tarnal, V., Picton, P., Harris, R. E., & Mashour, G. A. (2018). Subanaesthetic ketamine and altered states of consciousness in humans. British Journal of Anaesthesia, 121(1), 249–259. https://doi.org/10.1016/j.bja.2018.03.011

Vos de Wael, R., Benkarim, O., Paquola, C., Lariviere, S., Royer, J., Tavakol, S., Xu, T., Hong, S.-J., Langs, G., Valk, S., Misic, B., Milham, M., Margulies, D., Smallwood, J., & Bernhardt, B. C. (2020). BrainSpace: a toolbox for the analysis of macroscale gradients in neuroimaging and connectomics datasets. Communications Biology, 3(1), 103. https://doi.org/10.1038/s42003-020-0794-7

Vrijdag, X. C. E., van Waart, H., Mitchell, S. J., & Sleigh, J. W. (2021). An Electroencephalogram Metric of Temporal Complexity Tracks Psychometric Impairment Caused by Low-dose Nitrous Oxide. Anesthesiology, 134(2), 202–218. https://doi.org/10.1097/ALN.0000000000003628

Yeo, B. T. T., Krienen, F. M., Sepulcre, J., Sabuncu, M. R., Lashkari, D., Hollinshead, M., Roffman, J. L., Smoller, J. W., Zöllei, L., Polimeni, J. R., Fischl, B., Liu, H., & Buckner, R. L. (2011). The organization of the human cerebral cortex estimated by intrinsic functional connectivity. Journal of Neurophysiology, 106(3), 1125–1165. https://doi.org/10.1152/jn.00338.2011

